# Human Engineered Heart Tissue Models for Pharmacological Studies-Inhibition of Small-Conductance Ca^2+^-activated K^+^ (K_Ca_2) channel

**DOI:** 10.1101/2025.09.30.679523

**Authors:** Agnieszka Nadel, Ewelina Kałużna, Agnieszka Zimna, Anna Kowala, Agata Barszcz, Urszula Mackiewicz, Natalia Rozwadowska, Tomasz Kolanowski

**Affiliations:** Department of Molecular Pathology, Institute of Human Genetics PAS, Poznan, Poland; Department of Clinical Physiology, Medical Centre of Postgraduate Education, Warsaw, Poland; Department of Molecular Pathology, Institute of Human Genetics PAS, Strzeszynska St 32, 60-47 Poznan, Poland

**Keywords:** SK channels, Ca^2+^-activated K^+^channels, atrial fibrillation, engineered heart tissues (EHT), iPSC-CMs, chamber-specific heart tissue models

## Abstract

Atrial fibrillation (AF) is a common cardiac arrhythmia increasing stroke risk. Current treatments show limited effectiveness and can cause serious side effects. At the moment there is a lack of well-established human models recapitulating complex cardiac microenvironment for studying potential AF therapies. To address the limitations of existing models we developed 3D in vitro human Engineered Heart Tissue (EHT) model using atrial and ventricular human-induced pluripotent stem cell-derived cardiomyocytes (iPSC-CMs) with either human atrial fibroblasts (HAF) or human ventricular fibroblasts (HVF).These chamber-specific EHTs exhibit differences in gene expression, ion channels and physiological responses, enabling studying Frank-Starling mechanisms and Ca^2+^ response. Using K_Ca_2 channel inhibitor AP14145, we observed increased contraction force and prolonged relaxation time, suggesting negative modulation of K_Ca_2.2/K_Ca_2.3 channels and increased atrial-effective refractory period (AERP). The EHT model provides a valuable platform for examining K_Ca_2 channel inhibitors impact and evaluating chamber-specific drug effects in controlled *in vitro* settings.

## 1. INTRODUCTION

Cardiovascular diseases (CVDs) are considered the leading cause of death in developing countries among all ethnic groups, therefore prevention and treatment of CVDs are the utmost priorities ^1^. Patients are struggling most often with coronary heart disease. However, a significant proportion of the population have been diagnosed with life-threatening arrhythmias, including the most common sustained atrial fibrillation (AF). AF is characterized by a rapid and irregular rhythm of the atria leading to symptoms like palpitations or shortness of breath significantly reducing patient’s quality of life^2^. Treatment of AF focuses on several key strategies-like rhythm control, rate control, prevention of blood clot formation and management of stroke risk^4^. To obtain optimal therapeutic results, the standard treatment includes pharmacotherapy and sometimes surgical interventions-from less invasive electrical cardioversion through minimally invasive surgical ablation to Cox maze procedure^5^. However, managing far-reaching complications, such as heart failure and increased stroke risk, remain challenging for the clinicians^3^. Moreover, despite extensive research, complex mechanisms underlying AF remain incompletely understood, resulting in hindered treatment and prevention^5^. Ectopic source from the pulmonary veins (PV) is believed to be the most common trigger for AF and was identified in 94% of patients^6^. However, in 20% of patients, additional localizations of ectopic sources were identified: vena cava, the crista terminalis, the coronary sinus, the ligament of Marshall, and the inter-atrial septum^7^.

A major challenge in cardiovascular studies is the complexity of the heart, which is considered a highly specialized muscular organ divided into four functionally distinct chambers^8^. Myocytes from those chambers have divergent ultrastructure, gene expression patterns, and activity of ion channels located in the membrane^9^. Nonetheless, in the regulation of cardiac excitability and contractility, ions and their channels are the central players^10^. Ca^2+^ ions and their voltage-gated Ca^2+^ channels are crucial not only for initiating cardiac excitation-contraction coupling but also for activating multiple downstream signaling pathways in response to changes in membrane potentials, regardless of location. Consequently, changes in the channel’s functionality can have a significant effect on heart function. Small-Conductance Ca^2+^-activated K^+^ (SK) channels play a key role in AF, as they are triggered by Ca^2+^ impulses from the pulmonary veins^11^. It has been proven, in AF models, that SK channels are significantly upregulated in ventricular myocytes and in pulmonary veins during the heart failure ^11^.

Taking into account all the evidence, SK channels have become a promising target for managing AF. SK_Ca_ current is carried by three subtypes of SK channels: K_Ca_2.1, K_Ca_2.2, and K_Ca_2.3, which can be expressed in different parts of the human heart ^12^. Recent studies focus on the K_Ca_2.2 and K_Ca_2.3 subtypes, which are predominantly expressed in the human atria ^13^ and have been directly linked to the human AF.

SK channels are composed of six transmembrane regions (TMs) and a single pore loop and are solely activated or deactivated in response to the binding or release of Ca^2+^. Activation occurs when each calmodulin (CaM) molecule binds two Ca^2+^ ions during the Ca^2+^ level increases due to Ca^2+^ influx through L-type Ca^2+^ channels and sarcoplasmic reticulum Ca^2+^ release (the systolic phase)^14,15^. Activation of SK channels in cardiomyocytes leads to repolarization and shortening of the action potential duration (APD). SK channels work in a negative feedback system – when the APD is prolonged, it leads to an elongated Ca^2+^ wave, which enhances the activation of SK channels, thus contributing to the shortening of APD^16^.

To modulate the Ca^2+^-dependent SK channel activation, atrial-specific inhibitor AP14145 has been identified as a negative allosteric modulator of K_Ca_2.2 and K_Ca_2.3 subtypes^17^. Widely described in many animal models - mice^17^, rats^17^, guinea pigs^18^ and pigs^19^, AP14145 has been associated with increased contraction force and a prolonged relaxation time, potentially impacting cardiac function and offering insights into therapeutic possibilities, particularly in conditions like AF.

Atrial-specific inhibitor AP14145 was tested in established Engineered Heart Tissue (EHT) model based on human induced pluripotent stem cell-derived cardiomyocytes (iPSC-CMs). We have found that EHT model provide physiological information of the influence of AP14145 inhibitor differentiating atrial and ventricular tissues, including increase in atrial-effective refractory period (AERP). This EHT model aims to bridge the gap in human cardiac research, offering a platform to advance the development of new pharmacological treatments for heart conditions.

## 2. MATERIAL AND METHODOLOGY

### 2.1. Cell culture

#### Induced Pluripotent Stem Cells (iPSC)

In the study, two human iPSC lines - iWTD2.3 and iBM76.3, derived respectively from bone marrow cells and skin fibroblasts of healthy individuals, were used. Both cell lines were previously characterized ^20,21^. Human iPSC were cultured in Essential8 (E8) basal medium with E8 supplements (Thermo Fisher Scientific) on Geltrex-coated plates (17 μg/cm^2^, Thermo Fisher Scientific) and passaged using Versene® solution (0.53 mM EDTA; Lonza). For this study, at least three independent differentiations, using two different iPSC lines-iWTD2.3 and iBM76.3 were performed.

#### Cardiac fibroblast cell cultures (CFs)

Two fibroblast cell lines were used for the study, representing the population of atria and ventricles of the heart, respectively. Human Atrial Fibroblasts (HAF) gifted by Prof. Ali El-Armouche and Prof. Kaomei Guan from Technische Universität Dresden, Dresden Germany were described in Kϋnzel *et al.* ^22^. Commercially available Human Ventricular Fibroblasts (HVF) (Cat. 74038, Applied Biological Materials Inc.) were also used. Both lines were cultured on Geltrex-coated T75 flasks (17 μg/cm ^2^, Thermo Fisher Scientific) with Iscove medium (Thermo Fisher Scientific) supplemented with FBS, MEM-NEA, and insulin and passaged with Trypsin-EDTA (Thermo Fisher Scientific).

#### Directed differentiation of iPSC into CM subtypes - 2D cardiomyocytes culture

Cardiac differentiation of iPSC was performed via WNT signaling pathway modulation. In brief, differentiation was initiated at 90% confluence on Geltrex-coated 12 well plates. Medium E8 was changed to cardiac differentiation medium (CDM) composed of RPMI1640 with HEPES and GlutaMax (Thermo Fisher Scientific), supplemented with 0.5 mg/mL human recombinant albumin and 0.2 mg/mL-L - ascorbic acid 2-phosphate. The cells were sequentially treated for 48 h with 4 μM CHIR99021 (STEMCELL Technologies) and for a further 48 h - 5 μM IWP2 (Merck Millipore). On days 4 and 6, the medium was changed to CDM without small molecules. On the last day of the differentiation process - day 8, the medium was changed to the cardiac culture medium (CCM) composed of RPMI1640 with HEPES and GlutaMax and 2% B27 supplement (Thermo Fisher Scientific). From this day on, the CCM medium was replaced every two days. For atrial subtype differentiation, 1 μM RA (Sigma-Aldrich) was supplemented at days 3–6 during differentiation.

#### CM passage and selection

iPSC-derived CMs (iPSC-CMs) were digested around day 15 using collagenase II (Thermo Fisher Scientific) and 0.25% Trypsin/EDTA (Thermo Fisher Scientific). After centrifugation, cells were resuspended in a passaging medium composed of RPMI1640 with HEPES +GlutaMax (Thermo Fisher Scientific) supplemented with 20% fetal bovine serum (HyClone™) and 2 μM Thiazovivin (Sigma-Aldrich) and seeded in a density 750-850k cell/well (6-well Geltrex-coated plates). Around day 20-25, the metabolic CMs selection was performed using a cardio selection medium composed of RPMI 1640 without glucose (Thermo Fisher Scientific), 0.5 mg/mL human recombinant albumin, 0.2 mg/mL L-ascorbic acid 2-phosphate, and 4 mM lactate (Sigma-Aldrich). Selection was conducted for 4 consecutive days. After selection, iPSC-CMs were cultured in CCM for at least 40 days to be ready for EHT formation.

#### Engineered Heart Tissues (EHTs) generation - 3D cardiomyocyte culture

After iPSC differentiation into chamber-specific cardiomyocytes, chamber-specific EHT has been developed. A modified Tiburcy *et al*. ^23^ protocol has been used to introduce atrial and ventricular cardiac fibroblasts as a novelty factor to increase model specificity. EHTs were generated by combining iPSC-derived atrial/ventricular cardiomyocytes and cardiac atrial/ventricular fibroblasts in a 2,5:1 ratio (4.08×10^6^ iPSC-CMs to 1.7×10^6^ CFs), which was experimentally determined to be optimal to generate the highest number of stable tissue over time. The cell mixture was combined with type I Collagen Solution, 6 mg/mL (Advanced Biomatrix) in environment of 2% B27 supplement without insulin (Thermo Fisher Scientific), 0.5 M NaOH (Sigma) and 20 µl/mL penicillin/streptomycin (Thermo Fisher Scientific). The mixed cell suspension was transferred into circular casting molds, where it condensed for 4 days and was cultured in SFMM (Serum-free Maturation Medium) medium composed of Iscove Medium with Glutamax (Gibco), 1% MEM non-essential amino acids solution, 300 µM L-Ascorbic acid 2-phosphate sesquimagnesium salt hydrate (Sigma), 2% B27 Supplement without Insulin, 100 ng/mL IGF-1 (PeproTech), 5 ng/mL (PeproTech), 10 ng/mL FGF-2(PeproTech) and 5 ng/mL TGF-β (PeproTech). On day 5, EHTs were transferred onto the passive stretchers and cultured till the day of the measurements. After another 10 days of isometric exercises, on day 14 of tissue formation, measurements were performed in SFMM without TGF*-*β.

### 2.2. Atrial and Ventricular iPSC - CMs characteristics

#### Flow cytometry

iPSC-CMs were fixed for 10 min at RT using 4% paraformaldehyde (PFA; BiosterBio/Prospecta), permeabilized for 5 min/4°C with 0.1% Triton X-100 (Sigma). Fixed samples were incubated overnight with antibody against cardiac troponin T (cTNT; mouse, Thermo Fisher Scientific MS295PABX, 1:200) and K^+^voltage-gated channel subfamily A member 5 (KCNA5; rabbit, Alomone Labs APC-004, 1:100), resuspended in blocking buffer (0.5% Triton X-100, 1% BSA, 5% donkey/goat serum). The next day, cells were washed three times and incubated with secondary antibodies: Goat anti-mouse Alexa Fluor 594 (Thermo Fisher Scientific MS295PABX, 1:500) and Donkey anti-rabbit Alexa Fluor 488 (Thermo Fisher Scientific A-21206, 1:500) resuspended in the blocking buffer for 1 h at 4°C. Subsequently, cells were analyzed using the CytoFLEX flow cytometer (Beckman Coulter Life Sciences). Unstained and secondary-antibody-only stained cells were used as negative controls. At least 10,000 events were analyzed per sample. The results were analyzed using the Kaluza software.

#### Immunostaining

Cells were initially seeded on cover glasses with a density of 3 × 10^4^ cells/cover glass and incubated for 24 h in standard culture conditions to ensure complete attachment. The next day, cells were washed twice with a relaxation buffer (5 mM EGTA and 5 mM MgCl_2_ in DPBS) for 5 min at 37 °C and subsequently fixed in a 4% PFA solution at 4°C for 20 min.

Fixed slides were permeabilized using 1% Triton X-100, and then unspecific epitopes were blocked with 5% goat/donkey serum and Subsequently incubated overnight (4°C) with primary antibodies against cardiac troponin T (cTNT; mouse, Thermo Fisher Scientific MS295PABX, 1:200), α-actinin (mouse, Sigma A7811, 1:200), myosin light chain 2 ventricular (MLC2v; rabbit, Protein tech 10906-1-AP, 1:200) and myosin light chain 2 atrial (MLC2a; mouse, Synaptic System 311 011, 1:200). After thorough washing, secondary antibodies: Donkey anti-mouse Alexa Fluor 488 (BD Pharmingen 561495, 1:500), Goat anti-rabbit Alexa Fluor 594 (Thermo Fisher Scientific A-11012, 1:500), Goat anti-mouse Alexa Fluor 594 (Thermo Fisher Scientific A-11005, 1:500), were applied for 1 h at room temperature. Nuclei were counterstained with Fluoroshield™ DAPI (Sigma). Imaging was performed using a DMi8 confocal microscope (Leica). For quantitative volumetric analysis of the images, the Leica Application Suite X (LAS X) or Fiji (ImageJ) software was used.

#### Culture contraction analysis

All the recording of 2D cell culture and EHT beating patterns were performed by ZEN 2 software. The beating frequency of 2D iPSC-CM cultures was determined by MotionVektor, and for EHT-by counting spontaneous muscle contractions within 30 seconds of video recording.

#### Pharmacological studies

AP14145 (Tocris) was dissolved in DMSO (Sigma-Aldrich). Action potential recordings and force measurements in the AP14145 curve and calcium curve with the inhibitor were performed 5 min after the addition of the tested dose of drugs.

### 2.3. Force measurements of EHT

To perform force measurements, EHTs were placed on stainless steel hooks attached to a force-transducer and a length controller of the Myograph System (Danish Myo Technology A/S, Model840MD) perfused with Tyrode’s solution (containing 40 mM NaCl, 5.4 mM KCl, 1.8 mM CaCl_2_, 1 mM MgCl_2_, 10 mM HEPES, and 10 mM glucose; pH 7.4) and electrically field stimulated (0.5 Hz, 5ms, 10V). Both - the passive and active forces generated by the tissues were continuously measured during the whole procedure. First of all, initial length standardization was performed by sequential steps of stretching 0.1 mm up to lack of further increase in active force measurement. The determined passive force was the starting point for subsequent stages of measurements: 1) Ca^2+^-response curve (0.2 –3.4 mM), 2) AP14145 inhibitory influence (0.01 - 1000 nM), 3) β-adrenergic stimulation with isoprenaline (Sigma-Aldrich; 0.01 - 10 µM) influence. All measurements were performed to create minimum 7-point, semi-logarithmic response curves to comply with pharmacological standards. Force and length signals were digitally recorded and analyzed using LabChart Pro 8 software (ADInstruments) (Figure S1).

### 2.4. Molecular studies

#### RNA isolation and quantitative PCR

Pellets of either iPSC-CMs 2D cultures or HAF/HVF were snap-frozen in liquid nitrogen and stored at –80°C. Total RNA was isolated using acid guanidinium thiocyanate–phenol–chloroform extraction. EHTs-tissues after measurements were collected and stored in RNAlater™ Stabilization Solution (Thermo Fisher Scientific) at −20°C, to be isolated using Bead-beat Total RNA Mini kit (A&A Biotechnology). RNA concentrations and quality were measured using a NanodropOne spectrophotometer (Thermo Fisher Scientific). RNA was reverse transcribed into cDNA using 100 ng of total RNA using SuperScript™ IV Reverse Transcriptase kit (Thermo Fisher Scientific). Quantitative PCR was performed using PowerUp™ SYBR™ Green Master Mix and CFX96 Touch TM Real-Time PCR Detection System and analyzed by CFX Manager Software (all supplied by Bio-Rad). Negative controls, reference genes, and each cDNA sample in triplicates were amplified independently on the same plate. All results were normalized to glyceraldehyde-3-phosphate dehydrogenase (GAPDH), which was also used as an internal control. Primer sequences for real-time PCR are listed in Table S1.

#### Statistical analysis

All data were analyzed with GraphPad Prism 7 software using one-way ANOVA or Student’s t -tests depending on the experimental design. Data are presented with mean ± standard error of the mean (SEM). Experiments were performed in triplicates. Results were considered statistically significant when the p-value was < 0.05 (*p< 0.05; ** p< 0.01; ***p< 0.001; ns p> 0.05).

Experiments were performed using two different iPSC lines, iWTD2.3 and iBM76.3. Due to their distinct genetic backgrounds, results from these lines are presented together and denoted as N=2. The “n” value represents the total number of independent samples analyzed for each iPSC line (iWTD2.3 and iBM76.3).

## 3. RESULTS

### 3.1. iPSC-derived cardiomyocytes (CMs) are able to recapitulate chamber-specificity in 2D differentiation conditions

Both iPSC lines demonstrated the capacity to differentiate into atrial and ventricular cardiomyocytes according to the presented differentiation protocol (Figure 1A, adapted from Cyganek L. *et al.*^21^.).

**Figure 1.**
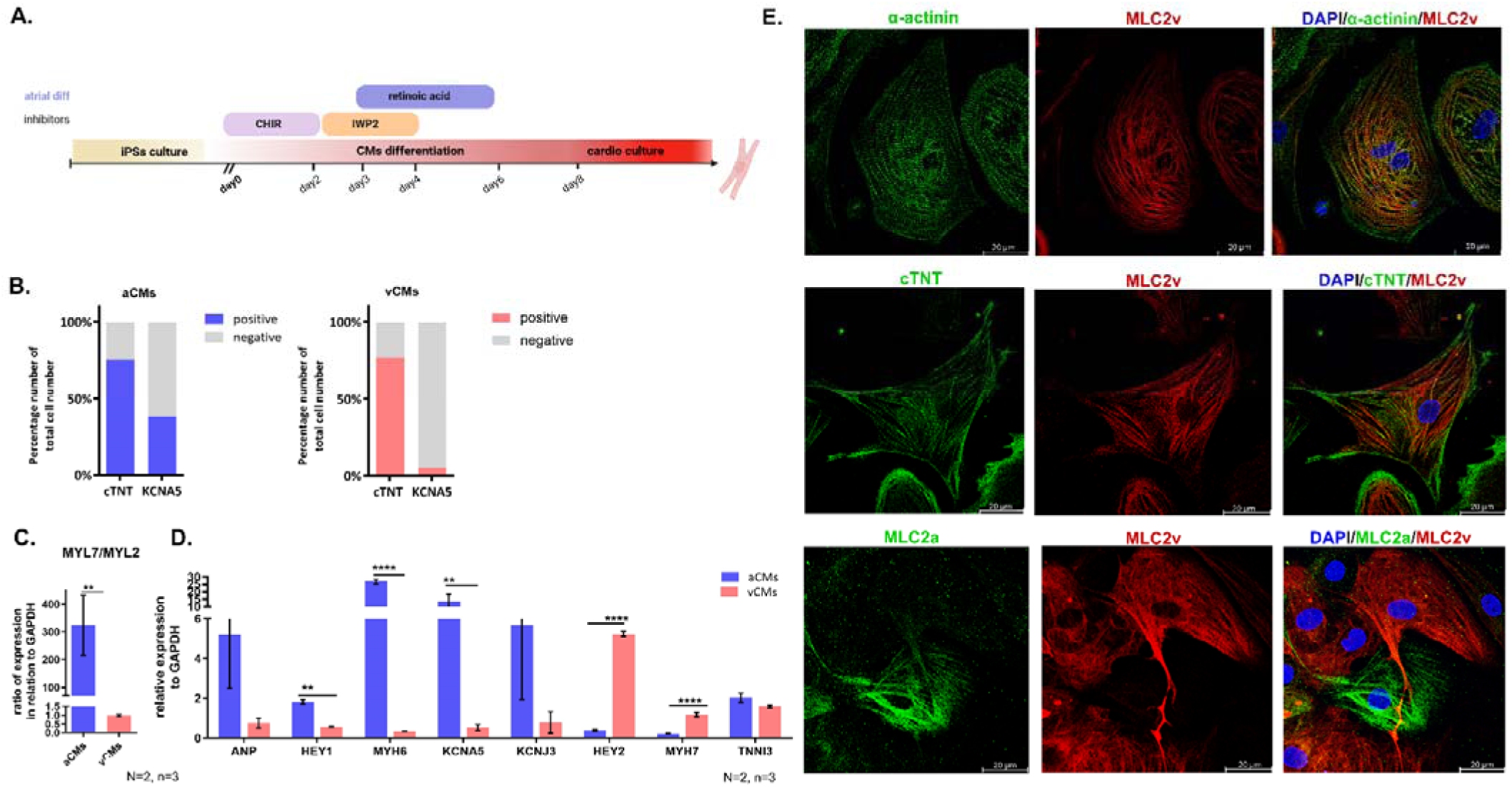
Chamber- specific iPSC-derived cardiomyocytes (CMs) A. A scheme of the CMs differentiation protocols. Small molecules, which modulate canonical WNT signaling - CHIR and IWP2, were applied to induce iPSC into mesoderm and to finally obtain ventricular cardiomyocytes (vCMs). Retinoic acid (RA) was used to induce the atrial cardiomyocytes (aCMs) differentiation. B. Flow cytometry analysis of iPSC-CMs subtypes- atrial/ventricular for cardiac troponin T (cTNT) and voltage-gated potassium channel (KCNA5). C-D. Real time PCR analysis. MYL7 (characteristic for aCMs) /MYL2 (characteristic for vCMs) ratio, which indicates the majority of CMs subtype in each population. Expression of of selected atrial (ANP, HEY1, KCNA5, KCNJ3, MYH6) and ventricular markers (HEY2, MYH7) in EHTs (3D) at day 14; normalized to GAPDH expression. D. Immunofluorescence staining of cardiomyocytes. Structural proteins, from the top: α-actin (green) with ventricular-type myosin light chain 2 (red); cardiac troponin T (green) with ventricular-type myosin light chain 2 (red); atrial-type myosin light chain 2 (green) with ventricular-type myosin light chain (red).. All values are expressed as mean ± SEM. *p<0.05, **p<0.01, ***p< 0.001, ****p≤ 0.0001 One-Way ANOVA was used for comparison.

Cytometric analysis of cardiac troponin type T (cTNT) and K^+^ Voltage-Gated Channel Subfamily A Member 5 (KCNA5) in cells differentiated with (RA+) or without (RA-) confirmed that the majority of cells in RA+ cultures expressed atrial marker (KCNA5), while ventricular phenotype (KNAC5 negative) was predominantly present in control RA-conditions (Figure S2). Only 8.43 % iPSC-CMs showed KCNA5 expression in RA-cultures, while treatment with 1 μM RA at days 3–6 generated 43,87% KCNA5^+^ CMs (Figure 1B).

To prove chamber-specificity on a molecular level, the expression of atrial and ventricular CM marker genes was analyzed using a real-time PCR technique. The atrial-specific markers included: structural - Myosin Heavy Chain 6 (*MYH6*), involved in cell signaling and gene regulation-Hairy and Enhancer of Split-Related Proteins 1 (*HEY1*) or connected with K^+^ channels: K^+^ Voltage-Gated Channel Subfamily A Member 5 (*KCNA5*) and Potassium Voltage-Gated Channel Subfamily J Member 3 (*KCNJ3*). The ventricular-specific markers CMs included: structural-Myosin Heavy Chain 7 (*MYH7*) and Hairy and Enhancer of Split-Related Proteins 2 (*HEY2*). Our data revealed that the *MYH6*, *HEY1* and *KCNA5* ratios were significantly increased in atrial CMs when compared to ventricular CMs*. MYH6* expression was 125 times higher inatrial CMs with p≤ 0.0001, HEY*1*-4 times with p<0.01 and *KCNA5*-30 times with p<0.01. Simultaneously, with 6 times higher expression of *MYH7* with a p≤ 0.0001 and 20 times higher expression of *HEY2* with a p≤0.0001 in ventricular CMs (Figure 1D). The *MYL7*/*MYL2* expression ratio (with *MYL7* encoding atrial and *MYL2* encoding ventricular myosin light chain 2) was 600 times higher in atrial CMs than in ventricular CMs (p < 0.01) (Figure 1C).

The atrial and ventricular subtype of 2D cardiomyocyte culture was also confirmed by immunofluorescent staining against myosin light chain 2: atrial (MLC2a) and ventricular (MLC2v) type (Figure 1E). Moreover, using antibodies against α-actin and troponin type T (cTNT) we were able to visualize the shape and components of cardiomyocyte’s contractility demonstrating cell structural integrity (Figure 1E).

### 3.2. Employing chamber-specific iPSC-CMs and fibroblasts in an EHT model provides high physiological specificity of the heart tissues

Current chamber-specific EHT models use chamber-specific iPSC-CMs combined with skin fibroblasts and extracellular matrix protein ^24^. However, these tissues lack full chamber-specificity, as skin fibroblasts are unable to transform into myofibroblasts; creating microenvironment that differs from *in vivo* conditions. To enhance tissue specificity, we provide chamber-specific cardiac fibroblasts-either HAFs or HVFs. This approach aim to create EHTs that more closely mimic the native cardiac tissue environment. The study design led to changes in physiological parameters, including contractility (Figure 2). Atrial cardiomyocytes exhibited a higher contraction rate, with an average of 111.25 ± 5.55 BPM, whereas ventricular subtypes contracted at a slower rate, with an average of 40.70 ± 5.17 BPM. This difference was statistically significant with a p<0.001 (Figure 2B). The addition of cardiac fibroblasts to the 3D model resulted in a reduction of the beat rate for both CMs subtypes. Transition from 2D monolayer to 3D EHT resulted in reduced beating frequencies for both cardiac types. The atrial CMs decreased from 115.11 ± 15.19 in monolayer culture to 62.20 ± 6.30 BPM in EHTs, while the ventricular CMs showed reduction from 40.00 ± 16.23 to 12.55 ± 1.01 BPM, at the same time.. The difference between chamber-specific EHT was statistically significant (p<0.001), reflecting the physiological differences observed between chambers in the human heart.

**Figure 2.**
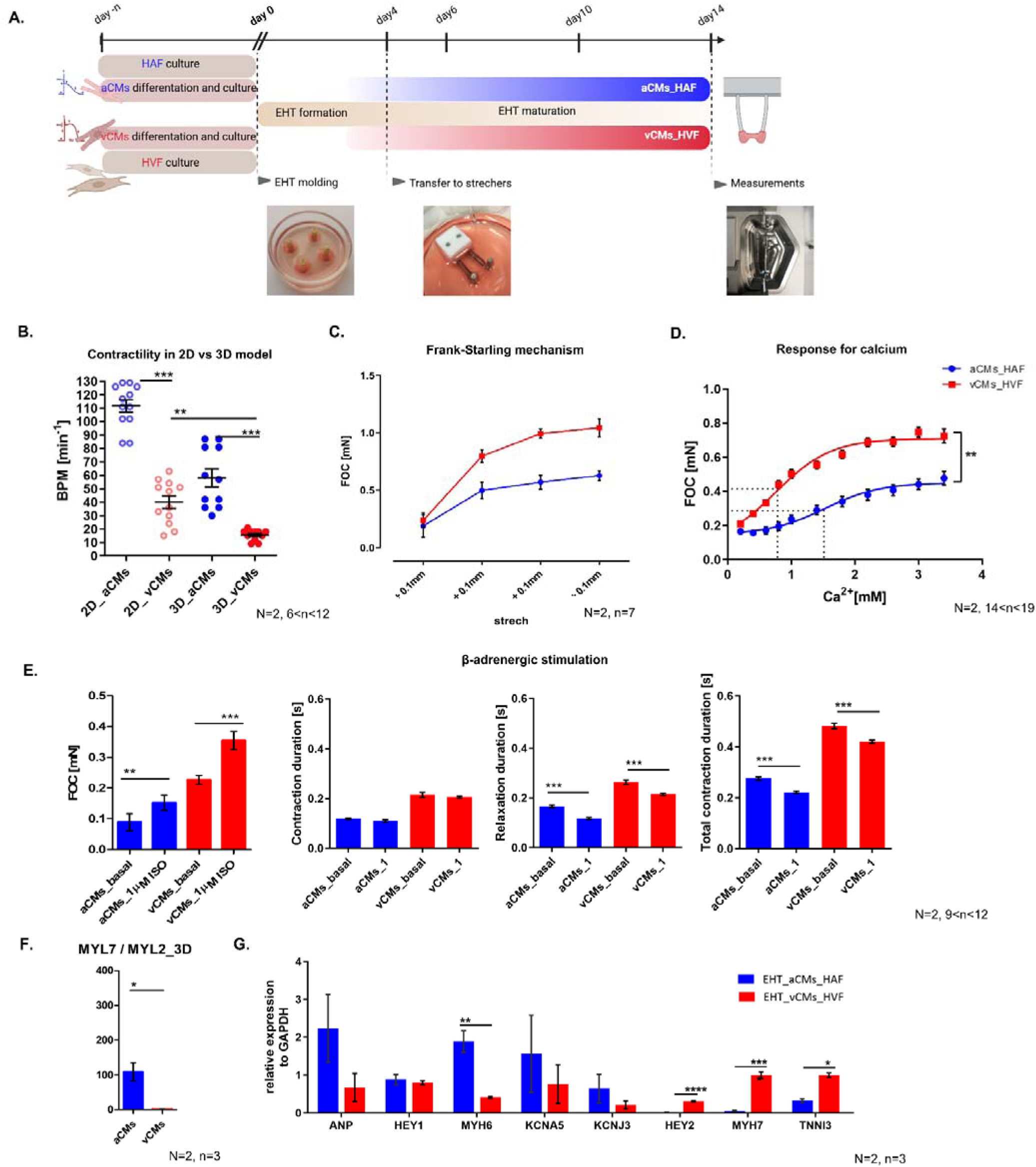
Chamber-specific EHT formulation. A. A scheme of chamber-specific EHT formation-by combining atrial cardiomyocytes with atrial cardiac fibroblasts and ventricular cardiomyocytes with ventricular cardiac fibroblasts, the establishment of chamber-specific tissue is possible. B. Contractility changes in atrial and ventricular cardiomyocytes in 2D/3D culture system. C. Differences between EHT subtypes in the Frank-Starling mechanism. D. Differences between EHT subtypes in the response for calcium. E. Differences between EHT subtypes in β-adrenergic stimulation. F-G. Real time PCR analysis. MYL7 (characteristic for aCMs) /MYL2 (characteristic for vCMs) ratio, which indicates the majority of CMs subtype in each population. Expression of selected atrial (ANP, HEY1, KCNA5, KCNJ3, MYH6) and ventricular markers (HEY2, MYH7) in EHTs (3D) at day 14; normalized to GAPDH expression. All values are expressed as mean ± SEM. *p<0.05, **p<0.01, ***p< 0.001, ****p≤ 0.0001; unpaired t-test has been used for comparison.

Measuring the force of EHT under different stretch-conditions, shows an increasing contraction force with greater stretch, which confirms Frank-Starling mechanism - the link between the initial length of myocardial fibers and the force generated by the cell. The force of contraction (FOC) within each EHT was quantified following incremental stretches of 0.01 mm in physiological calcium conditions of 1.4 mM. Starting from a pre-stretch baseline of 0.3 mN, FOC exhibited a logarithmic increase with each additional stretch until reaching a plateau phase after approximately the fifth elongation (Figure 2C).

In the study, EHTs were subjected to increasing concentrations of Ca^2+^ in Tyrode solution, and the force of contraction was measured. Both atrial and ventricular tissues showed a rise in contraction force as Ca^2+^ levels increased (0.4mM – 3.4mM range). The half-maximal Ca^2+^ response (logEC_50_) values for the atrial EHT were 1.529 mM and 0.754 mM for ventricular EHT-indicating the potency. The results showed a sigmoidal relationship between force and Ca^2+^ concentration, with a maximal force at high Ca^2+^ concentrations (Figure 2D). This relationship was observed in both atrial and ventricular EHTs.

Beta-adrenergic stimulation using 1 µM isoprenaline resulted in an increase of FOC in both atrial and ventricular EHT. Atrial FOC increased by 1.15 times compared to the basal state, with a p-<0.01. Ventricular tissue responded 1.7 times higher compared to the basal state (p-<0.001). Moreover, isoprenaline also affected contraction duration. Atrial contractions became shorter, decreasing from 0.27 to 0.21 s of total contraction duration time (tCD), while ventricular contractions became significantly shorter, decreasing from 0.48 to 0.42 s (, p<0.001) for tCD (Figure 2E).

To confirm the atrial-specificity of the model, the expression of atrial-like markers was confirmed by performing the real-time PCR. . It was found that all atrial EHTs had higher expression of atrial-specific markers compared to ventricular EHTs: structural-Myosin Heavy Chain 7 (*MYH7*); K^+^ ion channels-Voltage-Gated Channel Subfamily A Member 5 (*KCNA5*) and Potassium Voltage-Gated Channel Subfamily J Member 3 (*KCNJ3*); and metabolic-atrial natriuretic peptide (*ANP*) than ventricular EHTs (Figure 2D-E). The *MYL7*/*MYL2* expression ratio (with *MYL7* encoding atrial and *MYL2* encoding ventricular myosin light chain 2) was 100 times higher in atrial EHT than in ventricular EHT (p < 0.05) (Figure 1C).

All presented results from the physiological investigation and molecular analysis demonstrate that EHT can be used to create models of different types of heart tissue that mimic *in vivo* situations. We managed to increase the physiological relevancy of atrial and ventricular EHT compared to generally accepted 3D models.

### 3.3. Atrial EHTs are confirmed to express higher levels of K_Ca_2.3 channels, as observed in human atrial tissue

AP14145 is a negative allosteric modulator that, by binding to selective SK channel subtypes K_Ca_2.2 and K_Ca_2.3, makes calmodulin less sensitive to Ca^2+^, thereby increasing the AERP (Figure 3A).

**Figure 3.**
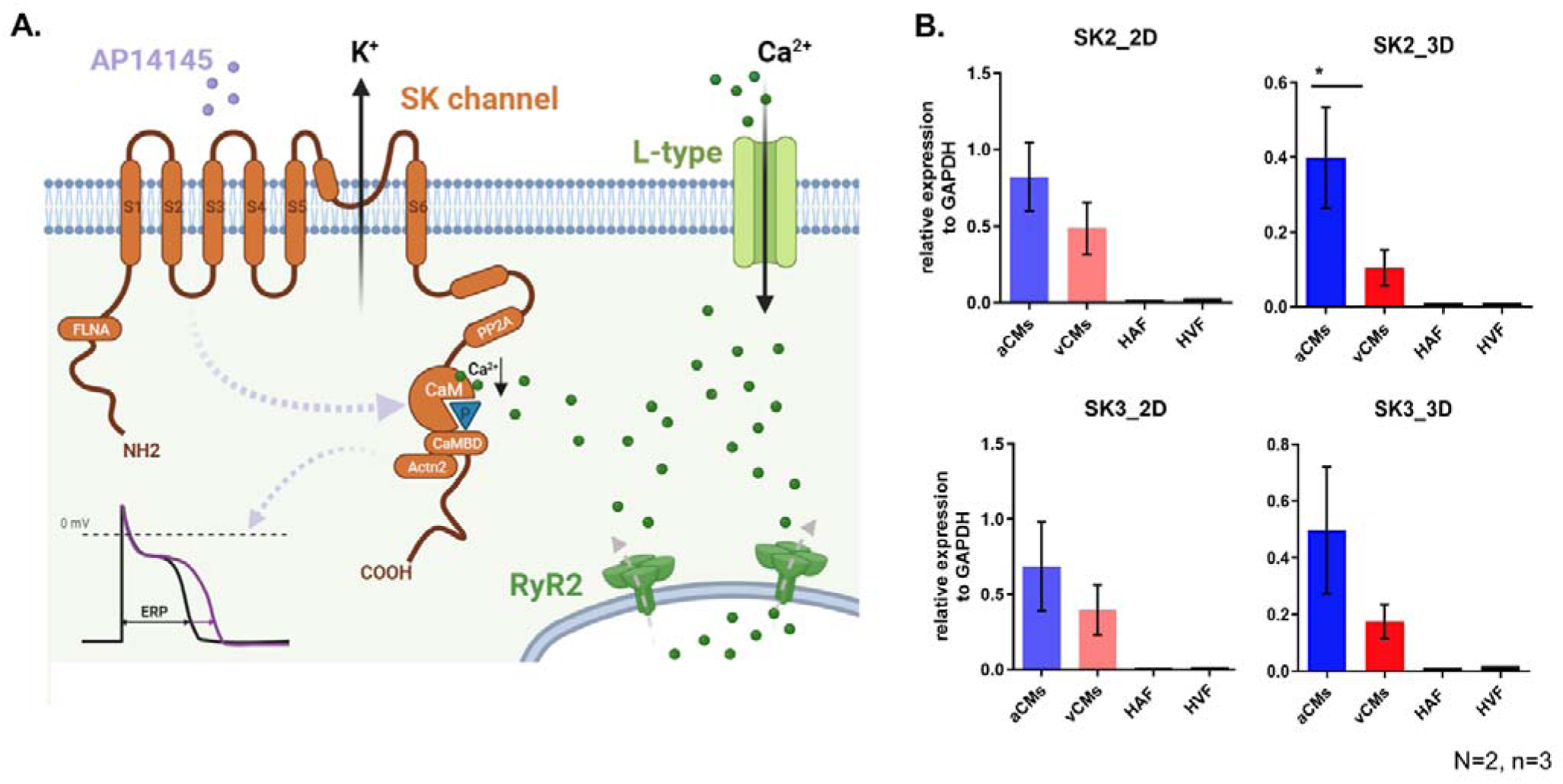
Mechanism of AP14145 influence on SK channels. A. Inhibition of small conductance Ca^2+^-activated K^+^ (K_Ca_2) channels activated by an increase in the concentration of intracellular Ca^2+^-prolongs atrial effective refractory period (AERP). B. Real-time PCR of selected K_Ca_2 subtypes-SK2 and SK3 in 2D/3D culture; normalized to GAPDH expression. All values are expressed as mean ± SEM. *p<0.05, **p<0.01, ***p< 0.001, ****p≤ 0.0001; unpaired t-test has been used for comparison.

We confirmed elevated SK channel expression in atrial cardiomyocytes in both 2D and 3D cultures (Figure 3B). Due to AP14145 selectivity, we focused on two subtypes - SK2 and SK3, which are both present in atrial cardiomyocytes. We detected mRNA-level expression of both genes in atrial and ventricular CMs, with more pronounced SK expression in atrial EHTs. However, the statistical significance was confirmed only in the case of SK3 expression (Figure 3B). No SK2 or SK3 expression was detected in atrial or ventricular fibroblasts. These findings demonstrate the presence of molecular targets for the AP14145 inhibitor.

### 3.4. AP14145 inhibitor effect is observed in atrial but not in ventricular 3D EHTs

AP14145 inhibitor effect was not observed in 2D culture condition (Figure S3), therefore AP14145 addition was tested on 3D EHTs, which to some extent better resemble human heart physiology than 2D models. According to kinetic studies and literature, the AP14145 is described with an IC_50_ of 1.1 μM for K_Ca_2.2 (SK2) and K_Ca_2.3 (SK3) channels. The activity of the inhibitor is limited to the atrial tissue, therefore confirming the selective targeting of AP14145. The application of logarithmically increasing inhibitor concentration provoked an exponential prolongation of atrial EHT contraction duration. This was in contrast to ventricular EHT, where increasing AP14145 concentration did not affect the total duration time (Figure 4A).

**Figure 4.**
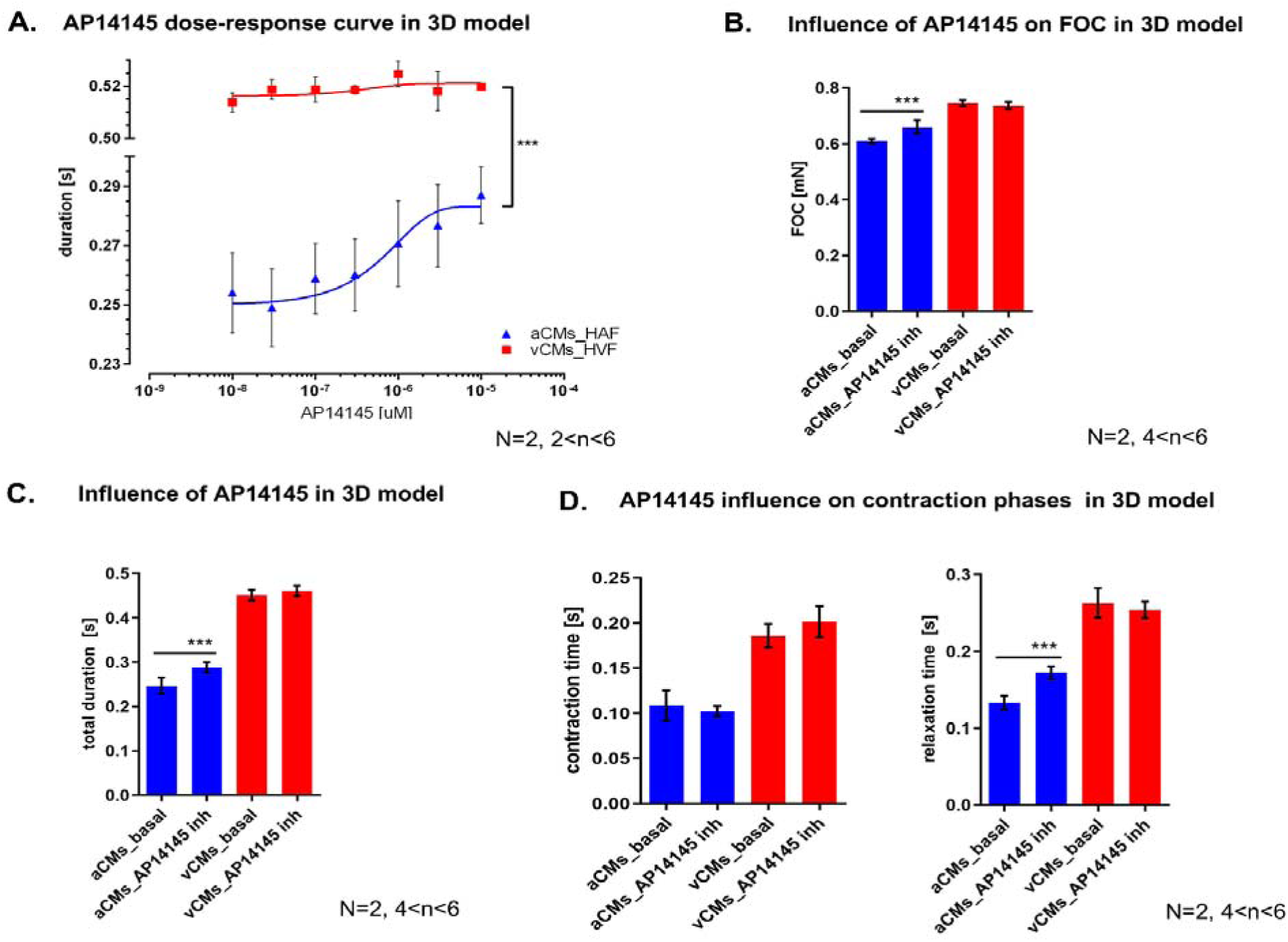
Influence of AP14145 on chamber-specific cardiomyocytes in 3D model. A. AP14145 response curve for atrial and ventricular EHT. B. Influence of 10 µM AP14145 on force of contraction (FOC) in atrial/ventricular EHT. C. Influence of 10 µM AP14145 on atrial/ventricular EHT contraction duration. D. Influence of 10 µM AP14145 on total contraction phases-contraction and relaxation time in atrial/ventricular EHT.

At the highest concentration tested on the AP14145 response curve (10 μM), we observed that atrial 3D models treated with the inhibitor showed enhanced FOC (Figure 4B). For atrial EHT, FOC significantly increased from 0.60 mN + 0.003 to 0.66 mN ± 0.008, with p< 0.001. In ventricular EHT, the FOC was maintained at the same level, approximately 0.73 ± 0.003 mN.

At this same concentration (10 μM), total contraction duration time also significantly prolonged in atrial EHTs-rising from 0.24 s ± 0.005 to 0.28 s ± 0.003 for atrial EHT, with p<0.001 (Figure 4C). In ventricular EHT, the duration time was maintained at the same level, (0.45 s ± 0.003), remaining stable and longer than atrial, thus maintaining physiological relevancy.

Moreover, a detailed analysis of contraction duration revealed that the influence of the inhibitor was restricted to relaxation time in atrial EHTs only, as SK2 and SK3 channels are active in the relevant phases (Figure 4D). Contraction time for atrial EHT was kept around 0.10 s ± 0.004 and for ventricular EHT-approximately, 0.19 s ± 0.004. At the same time, presenting the influence of 10 µM AP14145 on the relaxation time of atrial EHT. Due to the pharmacological incubation, relaxation time was prolonged from 0.13 s ± 0.002 to 0.17s ± 0.003, with p<0.001, indicating a statistically significant difference. The relaxation time for ventricular EHT was maintained at the same level, around 0.26 s ± 0.005 (Figure 4D).

## 4. DISCUSSION

The frequency of Atrial Fibrillation occurrence and its significant impact on patient’s well-being highlights the urgent need for finding a more effective treatment for AF. Exploring Small-Conductance Ca^2+^--Activated K^+^ (SK) channels as potential targets for AF treatment can provide promising, more effective and better-tolerated therapies. In this study, we shed additional light on the effects of the AP14145 selective SK-channels inhibitor. More importantly, we have created a fully atrial Engineered Heart Tissue (EHT) model and demonstrated its potential value in pharmacological efficiency evaluation of future therapies. While we did not directly assess pharmacological safety, the model’s ability to reflect known drug effects suggests its utility in early-stage safety screening. The presented 3D EHT model has the potential to match mature heart physiological properties, providing a valuable platform for studying the effects of drug candidates on cardiac function.

EHT model provides the unique opportunity to study physiological parameters of human heart tissue such as Frank-Starling mechanism, Ca^2+^-force relationship, and β-adrenergic stimulation using myographs and force transducers, which gives an opportunity to monitor the major parameters of heart function: force, and contraction/relaxation kinetics. Another physiological parameter possible to measure in the EHT model is the Ca^2+^-force relationship. When the concentration of Ca^2+^increases within the cytoplasm, it binds to troponin, shifting the position of tropomyosin and enabling myosin-actin binding. This leads to a stronger force of contraction. The third characteristic physiological parameter of the heart is the β-adrenergic stimulation during exposition to an adrenaline analog. To validate the β-adrenergic stimulation in EHT model, we used isoprenaline, a non-selective β1 and β2 adrenoceptor agonist. Isoprenaline increases heart rate and contractility by binding to β-adrenergic receptors in cardiomyocytes and enhancing Ca^2+^ influx and release from the sarcoplasmic reticulum. Consequently, the higher concentration of Ca^2+^ within the cell leads to more robust and more frequent heartbeats.

The introduction of the EHT model based on iPSC-CMs and cardiac fibroblasts represents a significant step forward in cardiac research. It allows for the investigation tailored to the chamber-specific tissue. The usage of human-derived EHT models offers many advantages, especially in terms of maturity, which allows us to study physiological parameters like the Frank-Starling mechanism^25^, responses to Ca^2+ 26^, and β-adrenergic stimulation^26^, which are not accessible in traditional 2D cultures (Figure 2C-E). Implementation of cardiac fibroblast and mechanical stimulation in EHT system enhances cardiomyocytes maturation, which is manifested by beating frequency ^27^. Cardiomyocytes cultured in the 3D model present a more rhythmic contraction pattern resembling that of *the in vivo* situation (Figure 2B)^8^. In the EHT model, mechanical communication may influence beating frequency, improving pacemaker cell integration. This process can lead to more mature ion channel expression patterns, which are essential for generating electrical signals and developing the contraction apparatus in cardiomyocytes^28^.

In our studies, we targeted SK ion channels, which are promising candidates for AF treatment. However, the development of new therapies is hampered by variable expression of SK2 and SK3 channel subtypes between species ^29^.

While SK2 and SK3 are often considered to be atrial-specific in humans ^13^, some studies suggest their presence also in ventricular tissue, although to a lesser extent— 33% lower in *KCNN2* and 50% lower in *KCNN3* ^30^. On the other hand, across various animal models, the expression of SK2 and SK3 channels varies with specificity in atrial and ventricular tissues. In mice, SK2 is predominantly expressed in atria, whereas SK3 is present in both atria and ventricles^31,32^. Rats exhibit SK2 and SK3 in ventricles, with SK2 also found in atria ^[33],[34]^. Guinea pigs show the occurrence of SK2 and SK3 in ventricular cardiomyocytes, along with consistent expression in atria and ventricles^17,35^. Pigs demonstrate expression of both SK2 and SK3 in both atria and ventricles^19^. Despite detected differences in the expression or main occurrence of SK2 and SK3, notably, the AP14145 prolongs the AERP without affecting the ventricles in most models, suggesting a specific impact on atrial electrophysiology across species^17–19^. Current animal studies suggest that AP14145 inhibitor may be a promising new antiarrhythmic drug for the treatment of AF. In the studies conducted thus far, AP14145 inhibitor has demonstrated efficacy in preventing shortening of the AERP and reducing the incidence of arrhythmias^17^. However, it’s important to acknowledge that some studies suggest potential ventricular effects, and further investigation is warranted to fully characterize the drug’s chamber selectivity.

In our studies, we observe that atrial EHT presents significant prolongation of total contraction duration, which may be caused by AP14145 influence on AERP (Figure 4C). Moreover, we detected the SK2 and SK3 expression in atrial and, to the lesser extent in ventricular EHT. These results are consistent with previous *in vivo* observations, which have demonstrated differential expression of these channels across cardiac tissues.(Figure 3B).

Given the established role of SK2 and SK3 channels in human atrial tissue, our investigation focused on the SK2/SK3-selective compound AP14145 - which effectively inhibited SK channel activity in aCMs in both 2D and 3D cultures. Applying AP14145 resulted in a prolonged contraction duration and relaxation phase (Figure 4A, 4C), confirming its role as a negative allosteric modulator of SK channels, as was presented by Smith *et al.* ^17^. Although we did not perform head-to-head comparative studies, our results are in line with published data, demonstrating AP14145 enhanced selectivity, potentially minimizing side effects, when compared to other SK channel blockers, such as NS8593 and AP30663, which reduce the sensitivity of CaM to intracellular Ca^2+ 36^.

Ca^2+^ plays two crucial roles in cardiomyocyte function: firstly, it directly increases contraction force, and secondly, it regulates SK channel activity by binding to calmodulin. During cardiomyocyte depolarization, Ca^2+^ influx into the sarcolemma triggers the release of additional Ca^2+^ from the sarcoplasmic reticulum storage, with high intracellular Ca^2+^ concentration promoting the interaction of actin and myosin filaments and leading to muscle contraction^37^. Thus, an increase in intracellular Ca^2+^ levels corresponds to a stronger contraction, which we also observed in our EHTs. Interestingly, the application of 10 µM AP14145, in the presence of stable 1.6 mM Ca^2+^ concentration showed an additional increase in FOC of atrial EHTs only (Figure 4EA). This phenomenon can be discussed from several aspects. First, AP1414, which triples the EC_50_ of Ca^2+^ on K_Ca_2.3 channels (from 0.36 to 1.2 μM)^17^, by reducing the sensitivity of CaM to intracellular Ca^2+ 38^ could lead to an increase in free-floating Ca^2+^ and indirectly influence the increase in FOC in atrial EHTs. SK2 and SK3 channels, are rather low abundance proteins, thus their quantitative influence on total Ca^2+^ availability in cardiomyocytes is questionable. Another more probable mechanism could rely on cell interdependence within the atrial tissue structure. Increased relaxation time upon AP14145 dosing may contribute to prolonged contraction, thus higher cumulative force being developed by the atrial muscle. While we hypothesize about the potential mechanisms behind the increased FOC, we acknowledge that we did not directly investigate the role of L-type Ca^2+^ currents or myofilament Ca^2+^ sensitivity. These aspects might be explored in more detail with the application of specific inhibitors or Ca^2+^ sensitizing compounds in subsequent investigations. It would be valuable to investigate whether similar increases in FOC have been reported in previous studies using AP14145 in atrial cardiomyocytes. Additionally, tracking Ca^2+^ transients in EHTs would provide important insights to confirm the hypothesis about changes caused by AP14145 treatment. Our study has several limitations. First, although the EHT tissue model has been significantly developed over the last few years, it still lacks full chamber-type purity. Although molecular and physiological experiments can clearly differentiate between atrial and ventricular tissues (Figure 2C), the current level of population purity used is not entirely homogeneous. There remains a degree of heterogeneity in the chamber-specific cardiomyocyte populations. Increasing purity at the cellular level will strengthen the model’s suitability as a valuable tool in pharmacological studies. Secondly, to get a better insight into the AP14145 inhibition mechanism, our studies would benefit from electrophysiological patch-clamp studies for tracking the activity of ion channels and may dispel current doubts regarding the influence of AP14145’s impact on AERP ^39^. Finally, two concepts require further elucidation: 1) AP14145’s influence on contraction apparatus-available Ca^2+^ levels, and 2) its inhibition of SK channels activity leading to prolonged relaxation and increased FOC. Wide range AP14145 and Ca^2+^ concentration studies in EHT’s measurements would help to evaluate these potential mechanisms. While these concepts offer a compelling mechanism, it is essential to conduct further experiments that consider the broader physiological context. To further strengthen the conclusions derived from our atrial 3D model, future experiments should include the implementation of other SK blockers, for instance apamin, as an additional comparator. Furthermore, it is important to acknowledge that our study was conducted in a control model. However, establishing the baseline effects of AP14145 on atrial electrophysiology in a healthy model is a crucial first step. Future research should aim to investigate the effects of AP14145 in an AF model to determine its therapeutic potential in the context of the disease and whether the mechanisms observed in the control model are maintained in the presence of AF-related remodeling. In summary, our study demonstrates the potential of the applications of EHT as a model for functional pharmacological investigations of new drug candidates. We have shown that the formation of EHT enhances the maturity of cardiomyocytes, modifying their beating patterns compared to traditional cultures.

Our findings reveal the specific effects of AP14145 on atrial EHT, resulting in prolongation of total duration time and increased FOC, while also raising questions about the mechanism of AP14145 Influence on the action potential and intracellular Ca^2+^-concentration. This investigation of the AP14145 inhibitor in EHTs underscores the value of the full chamber-specific engineered heart tissue model, which incorporates both atrial or ventricular cardiomyocytes and fibroblasts for enhanced functional pharmacological research. EHT model provides a more physiologically relevant and translational platform for drug discovery and development.

## CRediT authorship contribution statement

**Agnieszka Nadel**: Conceptualization, Investigation, Data curation, Visualization, Formal analysis, Methodology, Project administration, Writing - original draft, Writing - review & editing. **Ewelina Kalu**ż**na**: Investigation. **Agnieszka Zimna**: Writing - review & editing. **Anna Kowala**: Writing - review & editing. **Agata Barszcz**: Investigation. **Urszula Mackiewicz**: Supervision, Writing - review & editing. **Natalia Rozwadowska**: Supervision, Funding acquisition, Resources, Writing - review & editing. **Tomasz Kolanowski**: Methodology, Supervision, Formal analysis, Funding acquisition, Resources, Writing - review & editing.

## Supplemental Material

Table S1

Figure S1 – S3

