## Supplemental figures for "Human Engineered Heart Tissue Models for Pharmacological Studies-Inhibition of Small-Conductance Ca^2+^-activated K^+^ (K_Ca_2) channel"

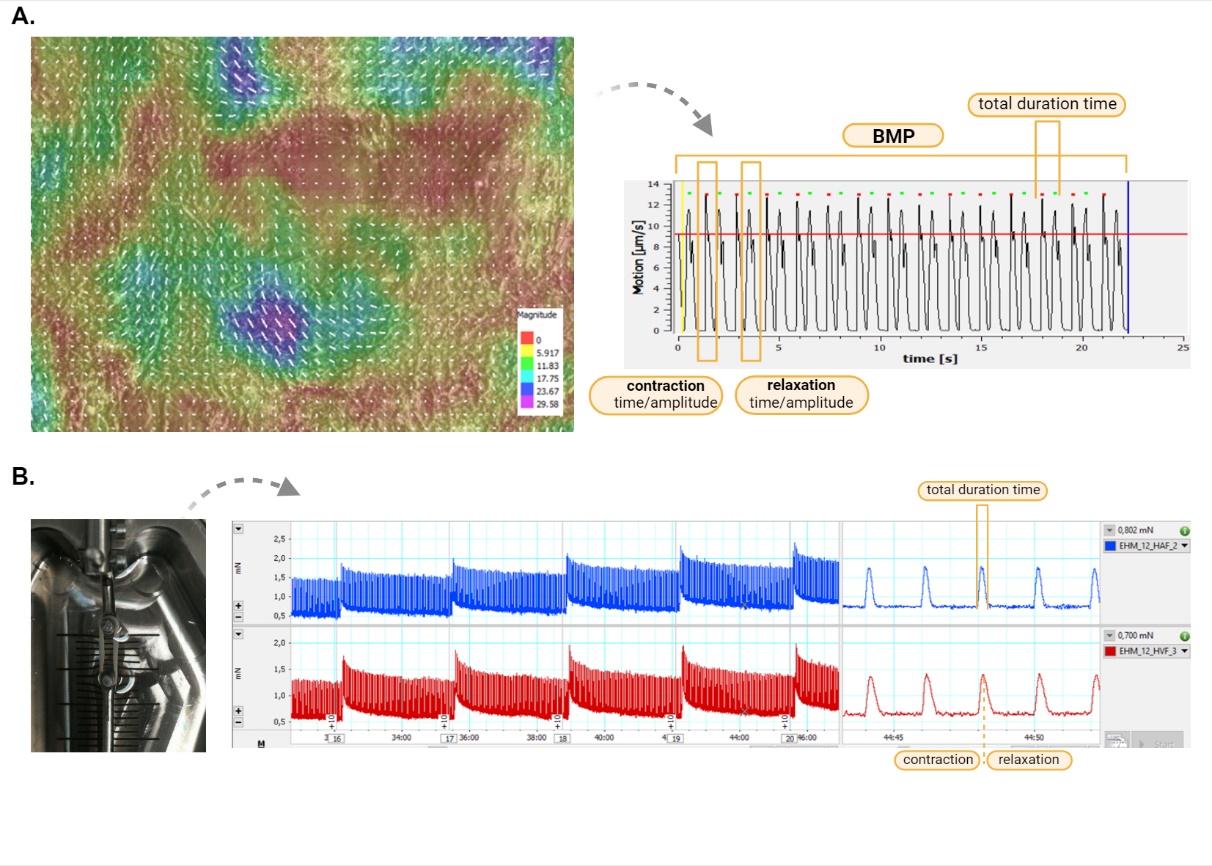

**Figure S1** An example of a processed movie. A. 2D ventricular CMs culture for contraction analysis in MotionVector software. B. 3D culture - aCM_HAF and CM_HVF in LabChart Pro 8 software.

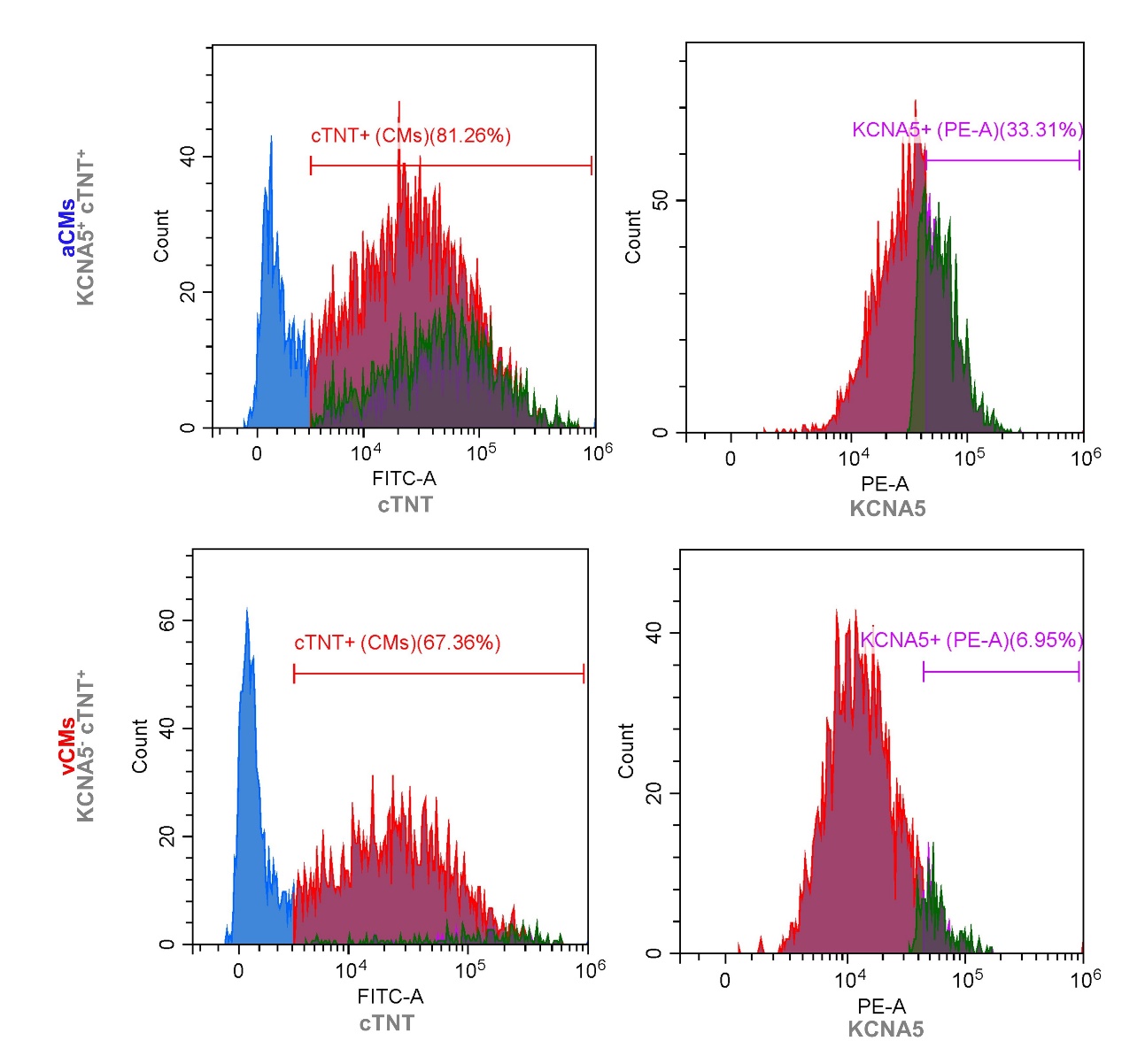

**Figure S2** Flow cytometry analysis of iPSC-CMs subtypes- atrial/ventricular for cardiac troponin T (cTNT) and voltage-gated potassium channel (KCNA5).

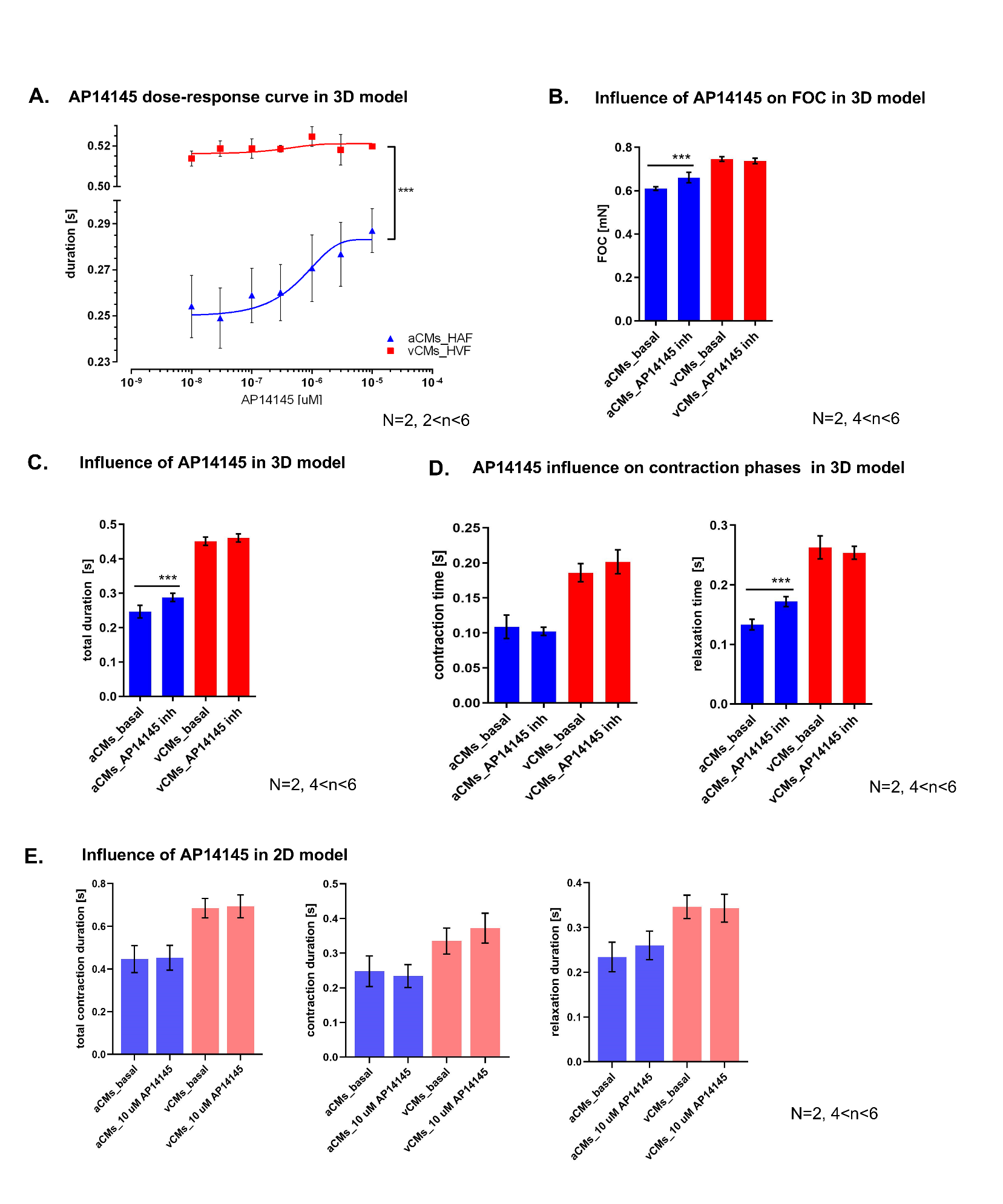
**Figure S3.** **AP14145 inhibitor effect is not observed in 2D culture condition.** 2D culture atrial and ventricular cardiomyocytes were treated with the highest tested concentration (10 µM) of AP14145. Within each cardiomyocyte subtype, the analysis of contraction patterns using MotionVector revealed no significant differences between the basal state and the highest dose of 10 µM AP14145 in 2D culture. The contraction duration for atrial CMs was approximately 0.31 s ± 0.035, while for ventricular CMs, it was 0.67 s± 0.007. At the same time, a detailed analysis of contraction duration revealed that treatment with 10 µM AP14145 did not affect the contraction or relaxation times of atrial and ventricular EHTs either.

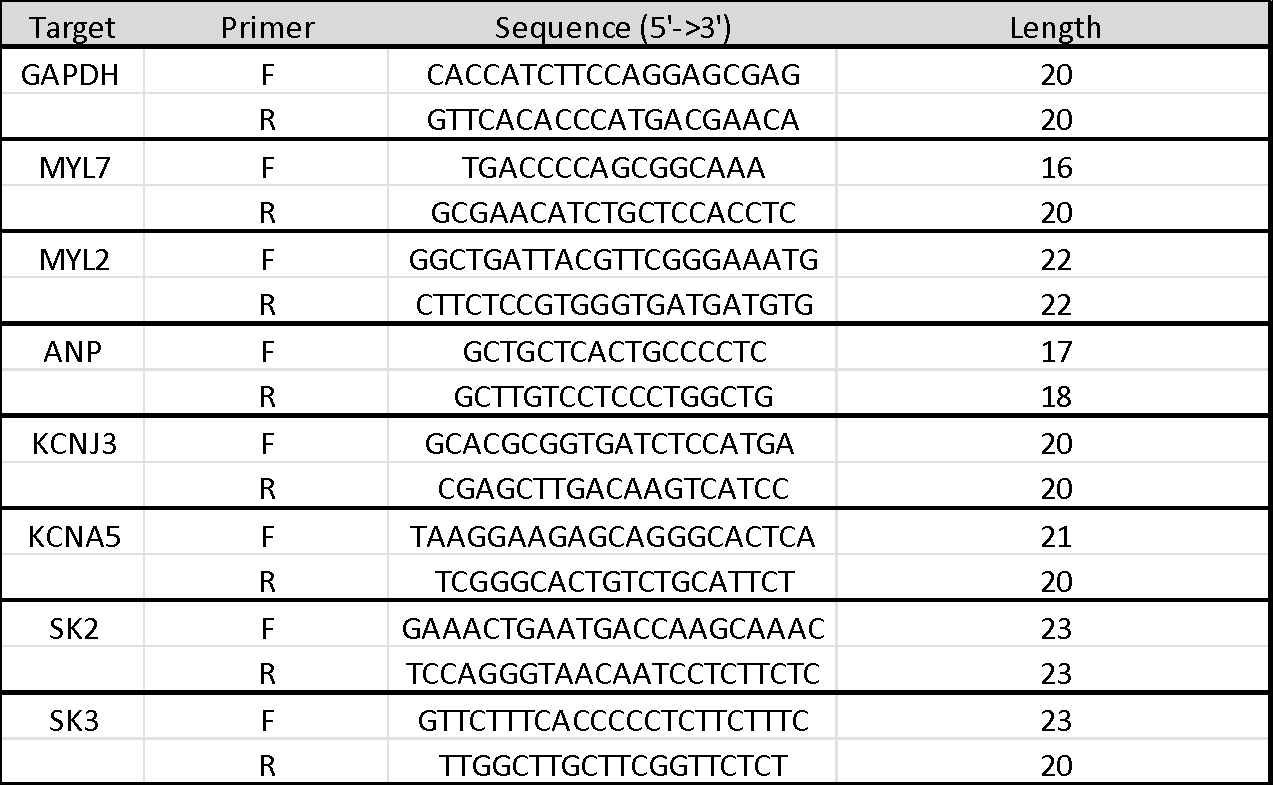

**Table S1** List of primers used during the molecular evaluation.
